# Genome expansion in early eukaryotes drove the transition from lateral gene transfer to meiotic sex

**DOI:** 10.1101/2020.03.21.001685

**Authors:** Marco Colnaghi, Nick Lane, Andrew Pomiankowski

**Affiliations:** CoMPLEX, University College London, London, United Kingdom, WC1E 6BT; Department of Genetics, Evolution and Environment, University College London, London, United Kingdom, WC1E 6BT

**Keywords:** Muller’s ratchet, Lateral gene transfer, bacterial transformation, sex and recombination, sexual reproduction

## Abstract

Prokaryotes generally reproduce clonally but can also acquire new genetic material via lateral gene transfer (LGT). Like sex, LGT can prevent the accumulation of deleterious mutations predicted by Muller’s ratchet for asexual populations. This similarity between sex and LGT raises the question why did eukaryotes abandon LGT in favor of sexual reproduction? Understanding the limitations of LGT provides insight into this evolutionary transition. We model the evolution of a haploid population undergoing LGT at a rate *λ* and subjected to a mutation rate *μ*. We take into account recombination length, *L*, and genome size, *g*, neglected by previous theoretical models. We confirm that LGT counters Muller’s ratchet by reducing the rate of fixation of deleterious mutations in small genomes. We then demonstrate that this beneficial effect declines rapidly with genome size. Populations with larger genomes are subjected to a faster rate of fixation of deleterious mutations and become more vulnerable to stochastic frequency fluctuations. Muller’s ratchet therefore generates a strong constraint on genome size. Importantly, we show that the degeneration of larger genomes can be resisted by increases in the recombination length, the average number of contiguous genes drawn from the environment for LGT. Large increases in genome size, as in early eukaryotes, are only possible as *L* reaches the same order of magnitude as *g*. This requirement for recombination across the whole genome can explain the strong selective pressure towards the evolution of sexual cell fusion and reciprocal recombination during early eukaryotic evolution – the origin of meiotic sex.

## INTRODUCTION

Understanding the origin and maintenance of sex in the face of multiple costs was long considered the ‘Queen of problems in evolutionary biology’ (Bell, 1982). Sexual reproduction breaks up advantageous combinations of alleles, halves the number of genes transmitted to the offspring, and is less efficient and energetically more costly than asexual reproduction (Bell, 1982; Otto and Lenormand, 2002; Otto, 2009). In spite of these disadvantages sex is a universal feature of eukaryotic life. The presence of common molecular machinery, widespread among all eukaryotic lineages, is a strong indication that the Last Eukaryotic Common Ancestor (LECA) was already a fully sexual organism (Schurko and Logsdon, 2008; Speijer et al., 2015). Meiotic genes are commonly found in putative asexual eukaryotes, including Amoebozoa (Lahr et al., 2011; Hofstatter et al., 2018), Diplomonads (Ramesh et al., 2005), Choanoflagellates (Carr et al., 2010) and even early diverging lineages such as *Trichonomas vaginalis* (Malik et al., 2008). Eukaryotic asexuality is not ancestral but a secondarily evolved state. The selective pressures that gave rise to the origin of meiotic sex must therefore be understood in the context of early eukaryotic evolution.

Phylogenomic analysis shows that eukaryotes arose from the endosymbiosis between an archaeal host and a bacterial endosymbiont, the ancestor of mitochondria (Williams et al., 2013; Martin et al., 2015; Zaremba-Niedzwiedzka, 2017). The presence of energy-producing endosymbionts allowed the first eukaryotes to escape the bioenergetic constraints that limit the genome size and cellular complexity of prokaryotes (Lane, 2014; Lane 2020). Extra energetic availability came with the evolutionary challenge of the coexistence of two different genomes within the same organism. As with other endosymbioses, the symbiont genome underwent a massive reduction, with the loss of many redundant gene functions (Timmis et al., 2004; López-Madrigal and Rosario, 2017). Alongside this, symbiont release of DNA into the host’s cytosol, caused the repeated transfer of genes to the host genome, many of which were retained, contributing to the massive genome size expansion during early eukaryotic evolution (Timmis et al., 2004; Martin and Koonin, 2006; Lane 2011).

Both the host and the endosymbiont, like modern archaea and proteobacteria, are likely to have been are capable of transformation – the uptake of exogenous DNA from the environment followed by homologous recombination (Bernstein and Bernstein, 2013; Vos et al., 2015; Ambur et al., 2016). This process involves the acquisition of foreign DNA, the recognition of homologous sequences and recombination, and therefore presents striking similarities with meiosis in Eukaryotes. The *Rad51*/*Dcm1* gene family, which plays a central role in meiosis, has high protein sequence similarity with *RecA*, which promotes homologous search and recombination in prokaryotes (Lin et al., 2006; Johnston et al., 2014;). It has been suggested that *RecA* was acquired by the archaeal ancestor of eukaryotes via endosymbiosis from its bacterial endosymbiont (Lin et al., 2006). Alternatively, the *Rad51*/*Dcm1* family could have evolved from archaeal homologs of *RadA* (Seitz et al., 1998). Regardless, the presence of this common molecular machinery and the striking similarities between these processes suggest that meiosis evolved from bacterial transformation (Schurko and Logsdon, 2008; Bernstein and Bernstein, 2013). But the selective pressures that determined this transition are still poorly understood.

Historically, the main focus of the literature on the origin and the maintenance of sex has been the comparison of sexual and clonal populations, or the spread of modifiers that increase the frequency of recombination (Bell, 1982; Otto, 2009). Recombination can eliminate the linkage between beneficial and deleterious alleles due to Hill-Robertson effects (Felsenstein and Yokohama, 1976; Barton and Otto, 2005), increase adaptability in rapidly changing (Hamilton, 1980; Gandon and Otto, 2007; Jokela et al., 2009) or spatially heterogeneous (Pylkov et al., 1998; Lenormand and Otto, 2000) environments, and prevent the accumulation of deleterious mutations due to drift predicted by Muller’s ratchet for asexual populations (Muller, 1968; Haigh, 1978;). These benefits of recombination outweigh the multiple costs of sexual reproduction and explain the rarity of asexual eukaryotes (Otto, 2009). But remarkably, they provide us with virtually no understanding of why bacterial transformation was abandoned in favour of reciprocal meiotic recombination. The real question is not why sex is better than clonal reproduction, but why did meiotic sex evolve from prokaryotic transformation?

Lateral Gene Transfer (LGT) has been recognised as a major force shaping prokaryotic genomes (Ochman et al., 2000; Lapierre et Gogarten, 2009; Vos et al., 2015). Unlike conjugation and transduction, mediated respectively by plasmids and phages, transformation is the only LGT mechanism to be exclusively encoded by genes present on prokaryotic chromosomes (Ambur et al., 2016), and is maintained by natural selection because it provides benefits analogous to those of sexual recombination (Levin and Cornejo, 2009; Wylie et al., 2010; Takeuchi et al., 2014; Vos et al., 2015). Recombination via transformation allows adaptation by breaking down disadvantageous combinations of alleles (Levin and Cornejo, 2009; Wylie et al., 2010) and preventing the fixation of deleterious mutations (Levin and Cornejo, 2009; Takeuchi et al., 2014). Some theoretical studies (Redfield et al., 1988; Redfield et al., 1997) suggest that transformation is only advantageous in presence of strong positive epistasis, a condition rarely met by extant prokaryotes. But more recent modelling work shows that transformation facilitates the elimination of deleterious mutations and prevents Muller’s ratchet (Levin and Cornejo, 2009; Takeuchi et al., 2014). As transformation provides similar advantages as meiotic sex, why did the first eukaryotes forsake one for the other? How did the unique conditions at the origin of eukaryotic life give rise to the selective pressures that determined this transition? In particular, is it possible that the massive expansion in genome size in early eukaryotes created the conditions for the evolution of a more systematic way of achieving recombination?

We know very little about the relation between genome size and the accumulation of deleterious mutations in populations undergoing transformation, as previous models either do not consider it explicitly (Levin and Cornejo, 2009; Wylie et al., 2010) or treat it as a constant parameter (Takeuchi et al., 2014). In order to evaluate the impact of genome size and recombination rate on the dynamics of accumulation of mutation, we develop a new theoretical and computational model of Muller’s ratchet in a population of haploid individuals, undergoing LGT at particular rates and subject to variable rates of mutation. Our model includes two new parameters, genome size and recombination length, which have not been taken into account by previous theoretical studies (Levin and Cornejo, 2009; Wylie et al., 2010; Takeuchi et al., 2014). We evaluate the effects of different selective landscapes, either uniform across the genome, or split between core and accessory genomes. The severity of the ratchet is evaluated using standard approaches for measuring the rate of fixation of deleterious mutations and the expected extinction time of the fittest class (Haigh 1978; Gordo and Charlesworth, 2000; Takeuchi et al., 2014). We suggest that systematic recombination across the entire bacterial genomes was a necessary development to preserve the integrity of the larger genomes that arose with the emergence of eukaryotes, giving a compelling explanation for the origin of meiotic sex.

## MATERIALS AND METHODS

We use a Fisher-Wright process with discrete generations to model the evolution of a population of *N* haploid individuals, subject to a rate of deleterious mutation *μ* per locus per generation, with LGT at a rate *λ*. The genome of an individual *j* is described by a state vector 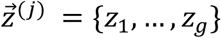, where *g* is the number of loci. Each locus *i* can accumulate a number of mutations {0,1,2….}. The components 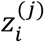 are the number of deleterious mutations at the *i*-th locus of the *j*-th individual. This allows us to keep track of the number of mutants in an individual and the distribution of mutations at each locus in the population. We define fixation of a mutant at a locus when the least-loaded class (LLC) at that locus is lost. As we neglect back-mutation, fixation of a mutant is permanent.

The genome-wide mutation rate *U* = *μg* is calculated as the product between the mutation rate per locus per generation and the number of loci (we assume that the mutation rate is constant across the whole genome). We introduce a new parameter *L*, the number of contiguous genes acquired with each LGT recombination event (i.e. the size of imported DNA), which has not been taken into account by previous theoretical studies (Levin and Cornejo, 2009; Wylie et al., 2010; Takeuchi et al., 2014). In order to avoid unnecessary complexity, we ignore the probability of ectopic recombination, and assume that DNA strands present in the environment (eDNA pool) are only stable for one generation before decaying irreversibly.

In the first part of this study, we assume that all mutations are mildly deleterious. Each mutation at a locus and across loci causes the same decrease in individual fitness *s*. Following previous studies of Muller’s ratchet (Haigh, 1978; Gordo and Charlesworth, 2000; Takeuchi et al., 2014), we choose a multiplicative function to model the fitness of an individual carrying *m* mutations, given by the formula *w*_*m*_ = (1 − *s*)^*m*^ (i.e. no epistasis). In the second part of the study we investigate more complex distributions of strength of selection across the genome. In particular, we differentiate between a strongly selected core genome and accessory genome under weaker selection. Which genes belong to the core and to the accessory genome is determined by random sampling using the MATLAB random number generator. The fitness of an individual that carries *m*_*i*_ mutations at locus *i* is given by 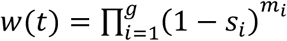, where *s*_*i*_ = 0.005 if locus *i* belongs to the core genome and *s*_*i*_ = 0.001 otherwise.

Each generation, the life history of the population follows the following processes. The new generation is obtained by sampling N individuals, with replacement, from the old population, using the MATLAB function randsample. The probability of reproduction is proportional to the individual fitness *w*_*m*_. The old generation dies, and their DNA forms the genetic pool from which the new generation acquires exogenous DNA (eDNA) for recombination. Each individual of the new generation acquires *n*^(*j*)^ new deleterious mutations, where *n*^(*j*)^ is a random variable drawn from a Poisson distribution with mean *U* = *μg*. The number of mutations and their position in the genome are determined using the MATLAB functions random and randi respectively. Each individual has a probability *λ* of undergoing LGT, determined by generating a random number using the MATLAB function rand. For each individual that undergoes LGT, a random donor is selected from the previous generation and *L* contiguous loci are randomly selected from its genome. The genome is assumed to be circular, so locus *g* is contiguous with locus 1. The corresponding components of the state vector of the recipient become equal to those of the donor. This can lead both to an increase or a decrease in the mutation load of the recipient. The process is started from a population free of mutation and repeated for 10,000 generations, with 50 replicates for a given set of parameter values.

Two measures *T*_*ext*,_ and *Δm*/*Δt* have been used to assess the effect of the ratchet (Haigh, 1978; Gordo and Charlesworth, 2000; Takeuchi et al., 2014). After recombination, we calculate the number of individuals in the least-loaded class (LLC). If a mutant reaches fixation at a particular locus, we mark this as *T*_*ext*,_ the time of extinction of the LLC. *T*_*ext*,_ gives an estimate of the time that a population can remain free of mutations. The second measure is the genome-wide rate of fixation *Δm*/*Δt*. This is calculated as the ratio between the total number of fixed mutations over the 10,000 generations of the simulation. The rate of fixation per single locus is the ratio between the genome-wide rate of fixation and genome size *g. Δm*/*Δt* is a measure of the rate of accumulation of mutations.

## Results

In absence of LGT, previous theoretical results (Haigh, 1978) have shown that, at equilibrium, the number of individuals in the least-loaded class (LLC) is *n*_0_ = *Ne*^.−*U*/S^. Without recombination and back-mutation, the loss of the LLC is an irreversible process – a “click” of the ratchet. The magnitude of *n*_0_ determines the likelihood that the least-loaded class becomes extinct because of stochastic fluctuations (i.e. genetic drift). High values of *n*_0_ increase the expected time of extinction of the LLC, whereas small values make the LLC more vulnerable to stochastic fluctuations (Muller, 1964; Haigh, 1978;). Therefore *n*_−_ is a good indication of the speed of the ratchet (Haigh, 1978). Expressing the genome wide mutation rate *U* as *μ* × *g* allows the equilibrium number of individuals in the LLC to be rewritten *n*_0_ = *Ne*^.−*g*/s^. The speed of the ratchet scales exponentially with genome size and mutation rate, and is negatively correlated with the strength of selection. Crucially, the impact of genome size is much stronger than that of population size (Figure 2). For example, a 2-fold increase in genome size can increase the speed of the ratchet by several orders of magnitude, whereas even a 10-fold reduction in population size has a rather meagre effect, except at low values (Figure 2). Therefore, any increase in genome size must be balanced by a proportional increase in strength of selection in order to avoid a drastic reduction of *n*_0_.

**Figure 1.**
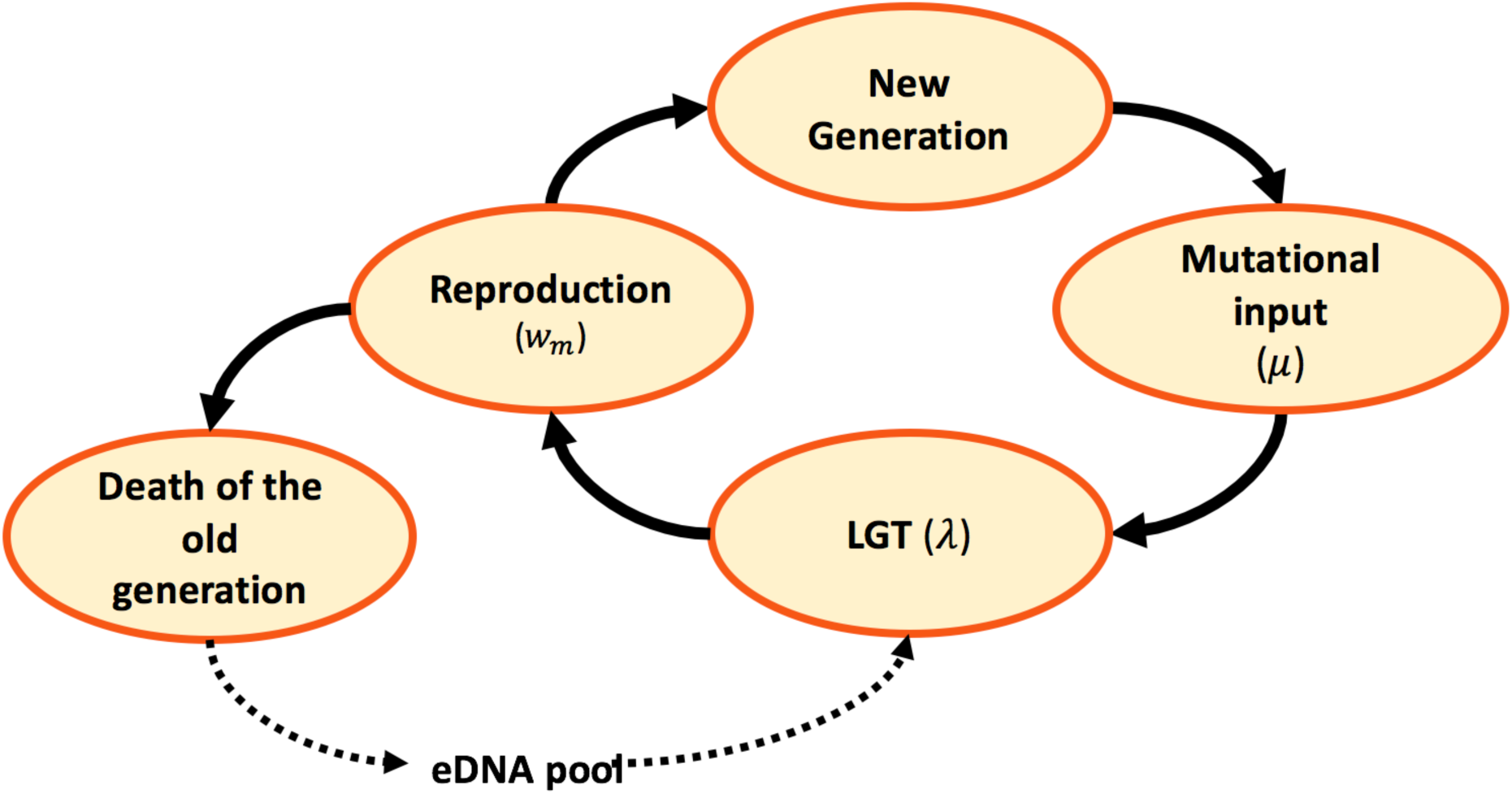
Model dynamics. After the birth of a new generation, new mutations are randomly introduced at a rate *μ* per locus. Following mutational input, eDNA is acquired from the environment and randomly recombined at a rate *λ* per genome. A new generation is then sampled at random, in proportion to reproductive fitness *w*_*m*_. The old generation dies and its DNA is released, constituting the eDNA pool for the new generation.

**Figure 2.**
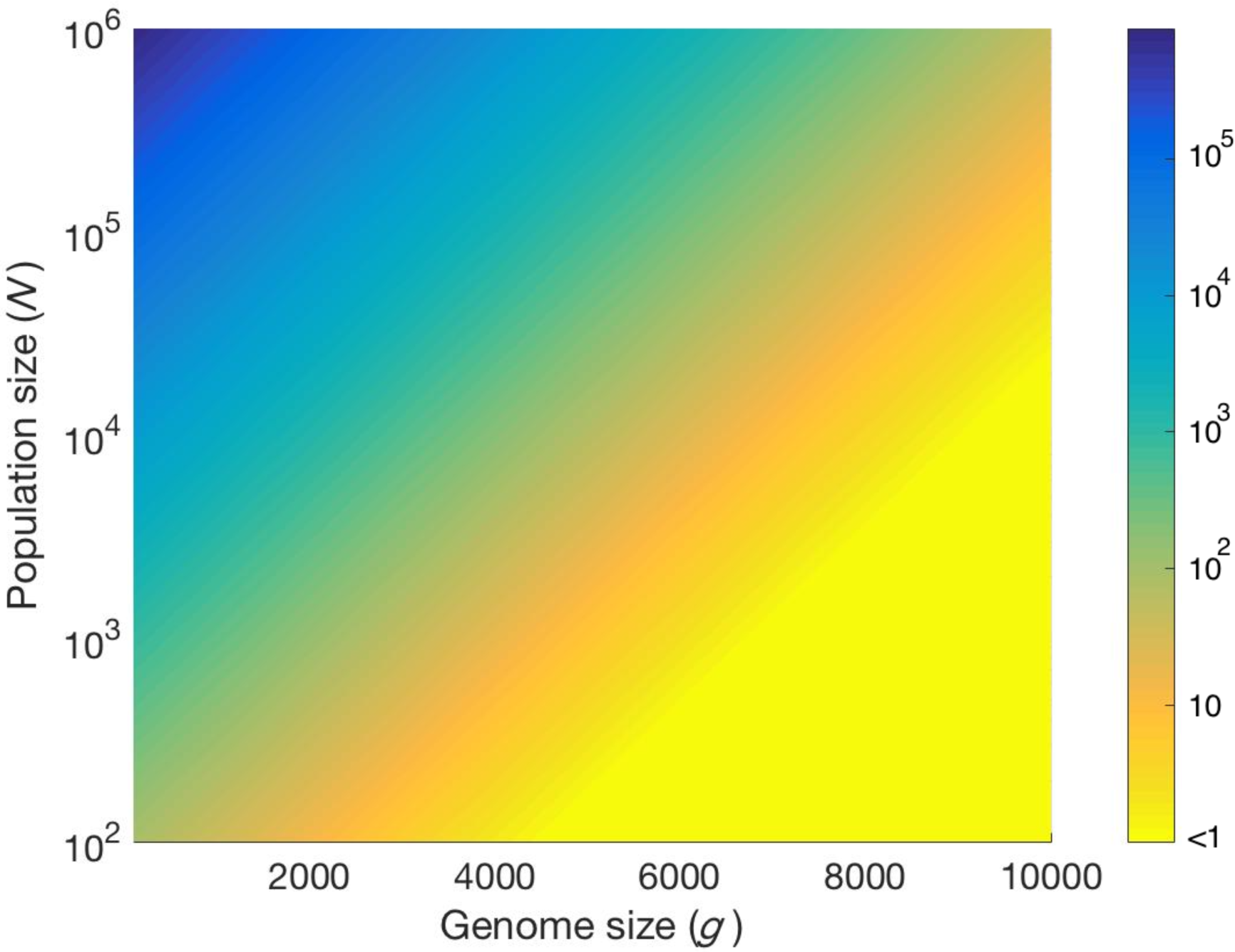
Genome size and population size determine *n*_0_. The equilibrium number of individuals in the least-loaded class (*n*_0_) is shown as a function of genome size (number of loci) and population size, with constant mutation rate *μ* = 10^−8^ and constant strength of selection *s* = 10^−3^.

### i. Uniform strength of selection

Genome size increases the severity of the ratchet, measured by *T*_*ext*,_, the expected extinction time of the LLC (Figure 3). Large genomes gain *de novo* mutations at a faster rate than small ones, leading to a decline in LLC extinction time, as there are more independent loci that can possibly fix for the mutant (Figure 3, no LGT). LGT reduces the severity of the ratchet and increase the expected LLC extinction time (Figure 3), making the population less vulnerable to stochastic fluctuations. The beneficial effect is more evident as recombination length (*L*) and LGT rate (*λ*) increase (Figure 3). However, as genome size (*g*) increases the expected extinction time plummets, rapidly approaching that of a clonal population with a larger genome, both in the presence of high (*λ* = 0.1, Figure 3A) and low (*λ* = 0.01, Figure 3B) LGT rates. The sole exception is when recombination length is of the same order as the magnitude of genome size (*L* = 0.2*g*, Figure 3). Only under this condition can increases in genome size be tolerated without a drastic decline in *T*_*ext*_.

**Figure 3.**
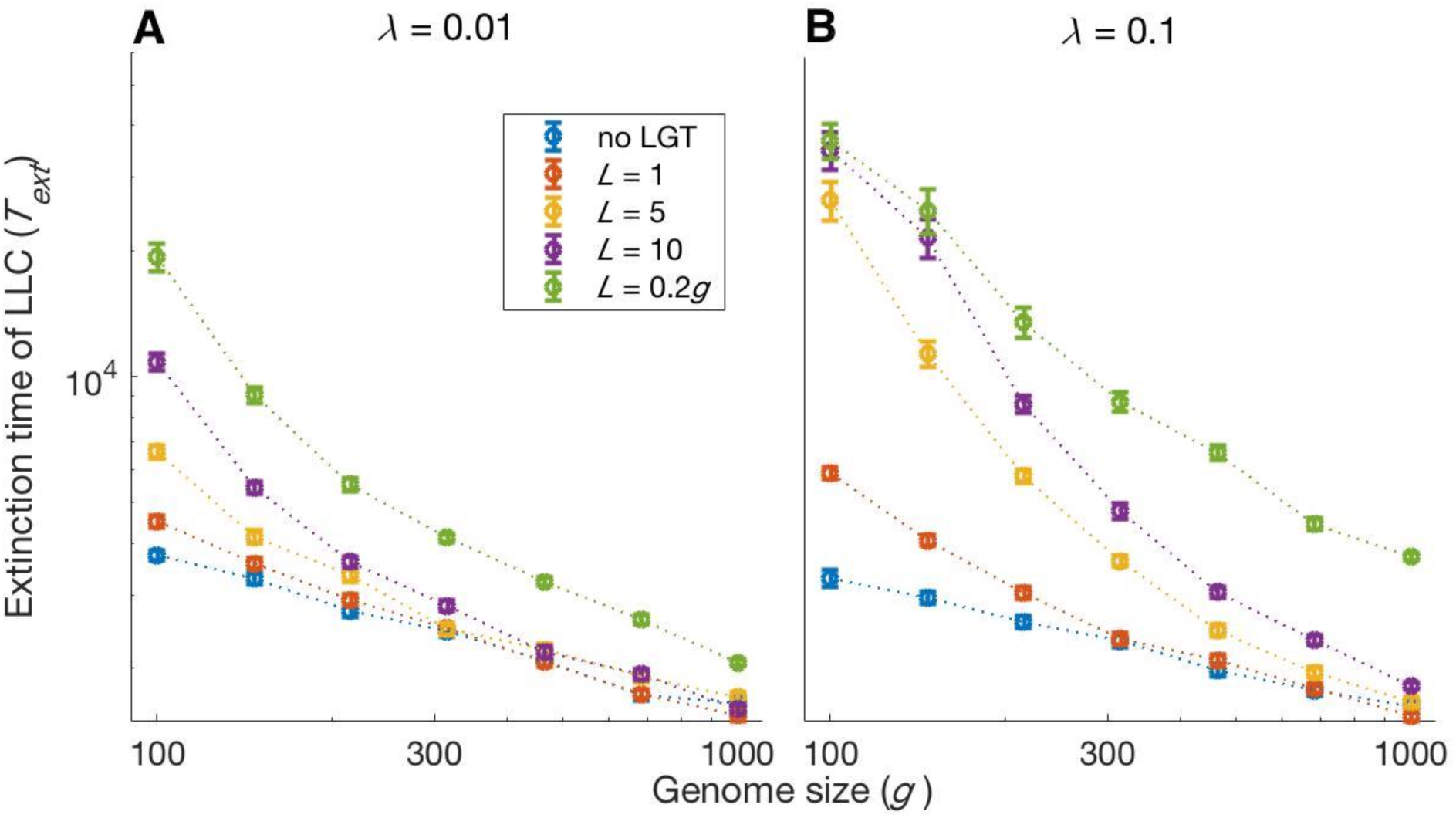
Impact of LGT and genome size on the ratchet. The average extinction time of the Least-Loaded Class is shown as a function of genome size (*g*) for various recombination lengths (*L*), in the presence of a) low (*λ* = 0.01) and b) high (*λ* = 0.1) LGT rates. The blue lines show the exinction time when there is no LGT, and is the same in a) and b. Parameters: *s* = 10^−3^, *N* = 5 × 10^3^, *μ* = 10^−7^. Error bars show the standard deviation over 50 independent iterations.

The rate of accumulation of deleterious mutations shows an analogous pattern (Figure 4). As genome size increases, the rate of mutation accumulation markedly increases, both genome wide and per locus (Figure 4). In a small genome, LGT reduces the speed at which mutations accumulate in a population, counteracting the ratchet effect both in the presence of a high (*λ* = 0.1) or low (*λ* = 0.01) LGT rate (Figure 4). This effect is more pronounced with higher LGT rates (*λ*) and longer recombination lengths (*L*). But even in presence of LGT, large genomes are subjected to higher rates of accumulation, comparable to those of a purely clonal population (Figure 4). Only when recombination length approaches the same order of magnitude as genome size (*L* = 0.2*g*) and occurs at high frequency (*λ* = 0.1) can LGT sufficiently repress mutation accumulation in large genomes (Figure 4B and 4D).

**Figure 4.**
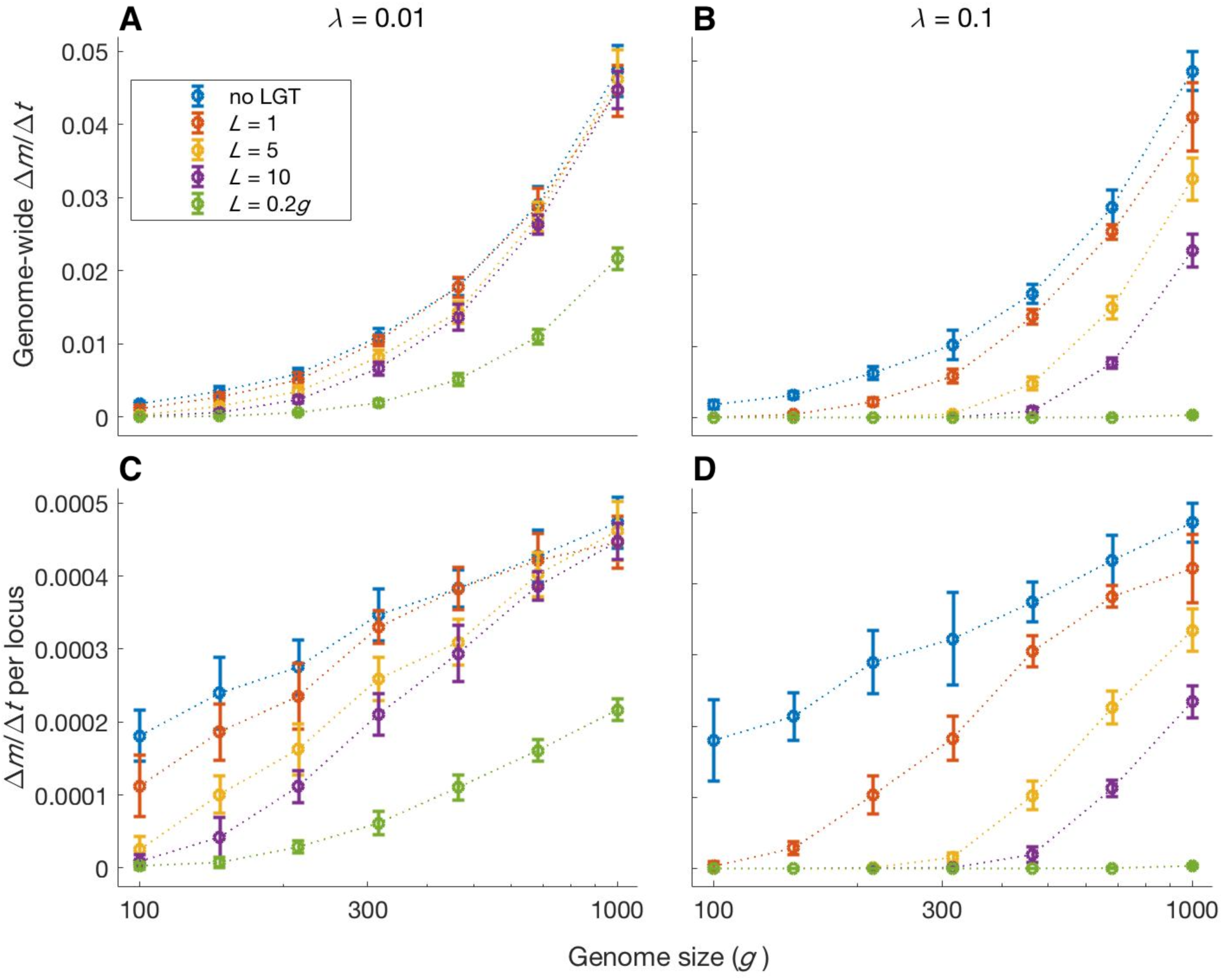
Impact of LGT and genome size on the rate of accumulation of mutation. The two upper panels show the mean genome-wide rate of fixation of deleterious mutations per generation, calculated over a time interval *t* = 10^5^ generations, as a function of genome size. Similarly, the two lower panels show the rate of fixation per single locus per generation. As genome size increases, LGT becomes less effective in reducing the mutational burden of a population. An increase in recombination length improves the efficiency of LGT in preventing the accumulation of mutations, but this beneficial effect declines rapidly with genome size. Only if recombination length is of the same order of magnitude as genome size (*L* = 0.2g) and the rate of LGT is high (*λ* = 10^−1^) large genomes can be maintained in a mutation-free state. Parameters: *λ* = 10^−2^ (left panels) and *λ* = 10^.−1^ (right panels), *s* = 10^−3^, *N* = 10^4^, *μ* = 10^−7^. Error bars show the standard deviation over 50 independent iterations.

### ii. Non-uniform strength of selection

Different loci in the genome are typically under different strengths of selection. In order to capture this fact in our model, we consider the core and accessory genomes differently. The size of the core genome is fixed (*g*_*c*_ = 50), while the accessory genome size varies as the genome expands. The core loci are under strong selection (*s* = 0.005) and the accessory loci are under weak selection (*s* = 0.001). Core and accessory loci are randomly distributed in the genome.

Under this selection regime, mutations preferentially accumulate in the accessory genome, where the strength of selection is lower, while the core genome accumulates mutations at a relatively slow rate (Fig. 5). Genome size expansion results in more severe ratchet effects, with a marked increase in the rate of mutations reaching fixation in the regions of the genome that are under weaker selection, alongside a moderate increase in core genome mutation fixation rate (Fig 5). LGT is effective in reducing the mutational burden, both in the accessory and in the core genome; but this beneficial effect is less evident in large genomes than in small ones (Fig 5). Recombination across the whole genome (*L* = 0.2*g*) completely eliminates fixation in the core genome, regardless of genome size, and markedly reduces the fixation rate in the accessory genome, facilitating genomic expansion (Fig 5).

**Figure 5.**
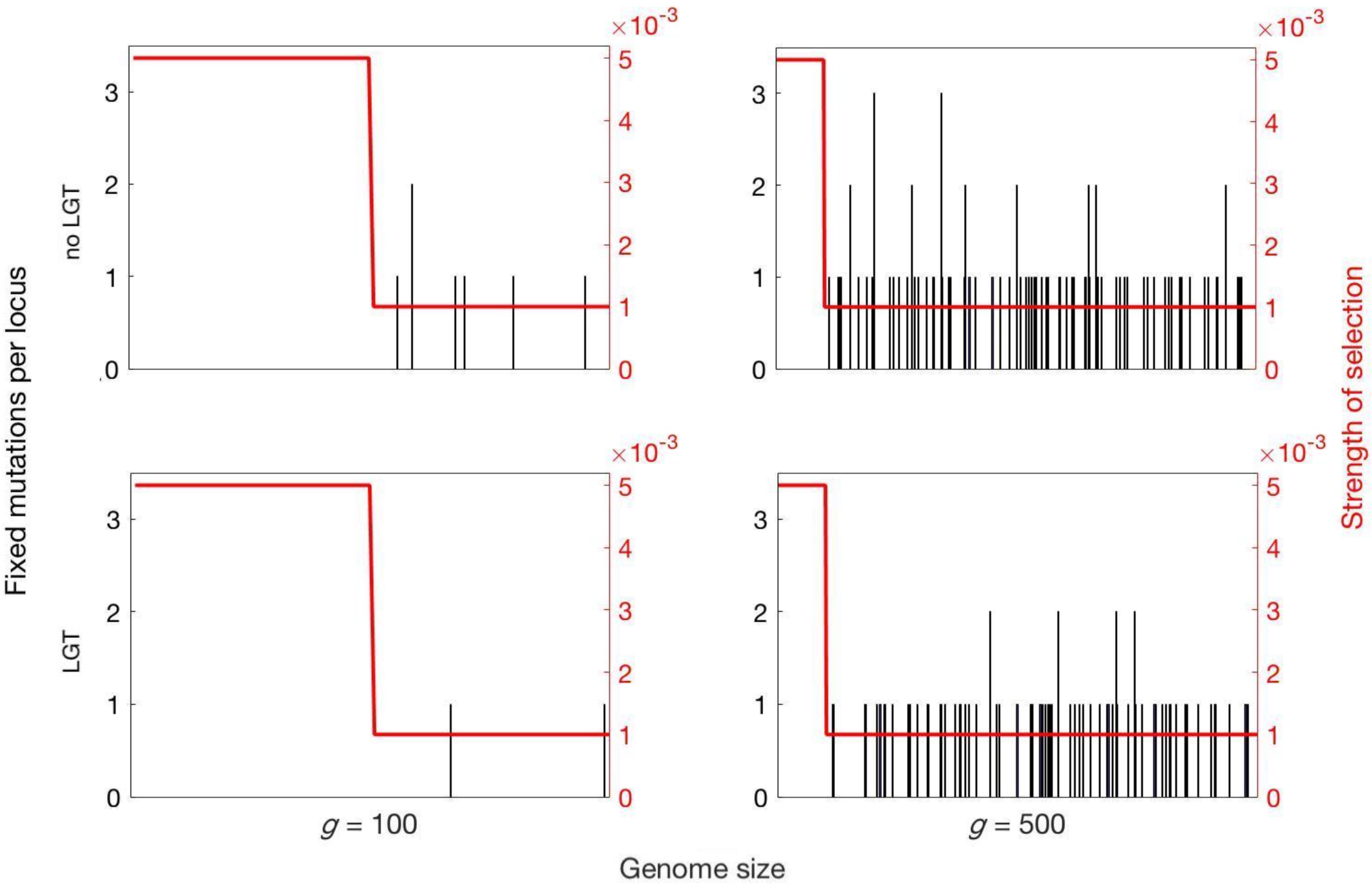
Fixation of mutations in the core and accessory genome. Fixed mutations in the core and accessory genome after *t* = 10^5^ generations for different genome size (*g* = 100, 500), without and with LGT (*λ* = 0.1, *L* = 5). Mutations preferentially accumulate in the accessory genome under weaker selection (s = 0.001), while the strongly selected core genome (*s* = 0.005) accumulates few or no mutations. The rate of fixation increases with genome size, while the benefits of LGT decline with genome size. Parameters: *λ* = 10^−1^, *L* = 5, *N* = 10^4^, *μ* = 10^.−7^.

**Figure 6.**
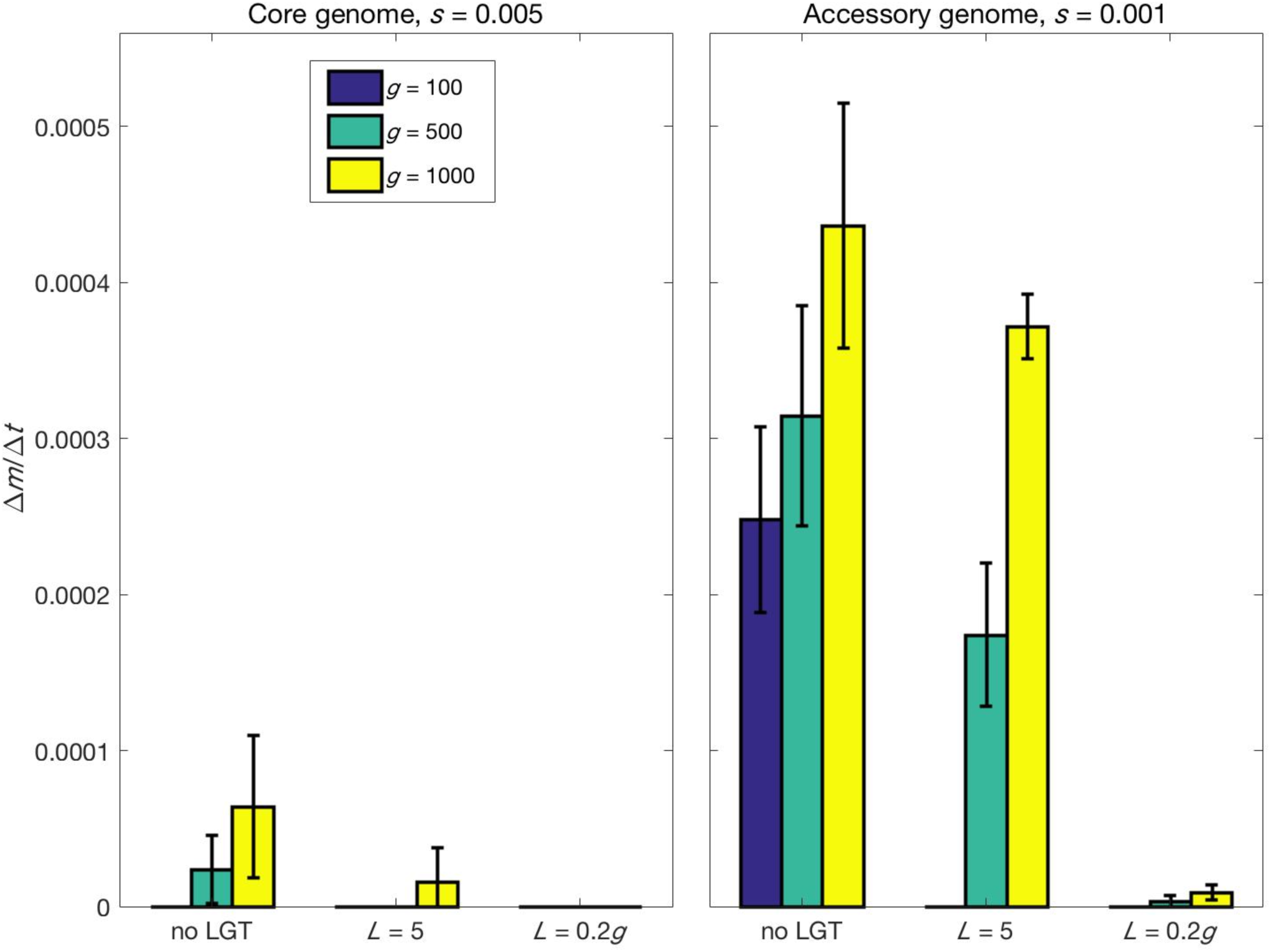
Rate of fixation of mutations in the core and accessory genome. The fixation of mutants in the core and in the accessory genome is shown after *t* = 10^5^ generations, normalised by genome size. A significantly higher number of mutations accumulate in those regions of the accessory genome that are under weaker selection. Genome size expansion increases the severity of the ratchet and the number of fixed mutations in the core and accessory genome. The introduction of LGT considerably reduces the mutational burden. Parameters: *λ* = 10^−1^, L = 5, *N* = 10^4^, *μ* = 10^−7^, *s*_*core*_ = 0.005 and *s*_*acc*_ = 0.001.

## Discussion

Asexual organisms are well known to be vulnerable to the effects of drift, which reduces the genetic variation within a population, causing the progressive and inescapable accumulation of deleterious mutations known as Muller’s ratchet (Muller, 1964; Haigh, 1978; Otto, 2009). In eukaryotes, sexual recombination counters the effects of genetic drift and restores genetic variance, increasing the effectiveness of purifying selection and preventing mutational meltdown (Otto, 2009). In prokaryotes, sexual fusion does not occur. But the exchange of genetic material does occur through transformation, the lateral gene transfer (LGT) and recombination of environmental DNA (eDNA). Meiotic recombination likely arose from bacterial transformation. Understanding the reasons why this transition occurred during early eukaryotic evolution are critical to a rigorous understanding of the Queen of problems in evolutionary biology, the origin of sex. Sex did not arise from cloning, as tacitly assumed in the classic theoretical literature, but from prokaryotic transformation, a very different question which we explored. To elucidate this transition, we examined the effectiveness of LGT at countering the dynamics of Muller’s ratchet, to understand where and why LGT becomes ineffective at maintaining genome integrity, necessitating the transition to sexual reproduction in early eukaryotes.

We assessed the effect of LGT on the severity of the ratchet using the expected extinction time of the least-loaded class and the rate of fixation of deleterious mutations. Unlike previous modelling work, we included genome size as a variable as opposed to a constant (e.g. 100 loci; Takeuchi et al., 2014). Genome size is plainly important in relation to the evolution of eukaryotes, which have expanded considerably in almost every measure of genome size (e.g. DNA content, number of protein-coding genes, size of genes, number of gene families, regulatory DNA content, Lane and Martin, 2010). Considering gene number in our model reveals a strong inverse relationship between genome size and the benefits of LGT. In small genomes, LGT is effective at preventing Muller’s ratchet, with long extinction times (Fig. 3) and low rates of mutation accumulation (Fig. 4), validating the results of previous theoretical studies (Takeuchi et al., 2014). However, we show that large genomes limit the efficiency of LGT and present a greater mutational target than smaller ones, increasing the overall input of mutations to the genome. This increases the severity of the ratchet leading to shorter extinction times and faster rates of mutation accumulation (Fig. 3-4).

The increased potency of the ratchet as genome size increases is ameliorated by an increase in the rate of LGT (*λ*; Fig. 3-4). Is this a viable option for prokaryotic species to enable them to expand genome size? In a number of species, LGT has been estimated as being the same magnitude (or higher) as the rate of nucleotide substitution, including *B. cereus* (Hao and Golding, 2006), *Streptococcus* (Marri et al., 2006), *Corynebacterium* (Marri et al., 2007), and *Pseudomonas syringae* (Nowell et al., 2014). Rates are highly variable among species (Croucher et al., 2012; Vos et al., 2015; Johnston et al., 2015). Competence for transformation can be induced by a range of environmental stressors including DNA damage, high cell density and limited nutrient availability (Bernstein and Bernstein, 2013). But LGT rates are constrained by eDNA availability, which depends on the amount of DNA in the environment and the degree of sequence homology (Croucher et al., 2012; Vos et al., 2015). The model predicts that higher LGT rates will strengthen purifying selection and favour the elimination of mutants. This result is compatible with the strong correlation observed between the number of horizontally transferred genes and genome size across a range of prokaryotes (Jain et al., 2003; Fuchsman et al., 2017). However, it is not clear to what extent the rate of LGT can be modified. Our modelling suggests that larger bacterial genomes are more likely to be sustained by higher rates of LGT, but the benefits of LGT as actually practiced by bacteria – the non-reciprocal uptake of small pieces of DNA comprising one or a few genes – are unlikely to sustain eukaryotic-sized genomes. In short, we show that LGT as actually practised by bacteria cannot prevent the degeneration of larger genomes.

Importantly, we show that the benefits of LGT also increase with recombination length (*L*; Fig. 3-4). In gram positive bacteria, recombination of large eDNA sequences is the exception rather than the rule (Croucher et al., 2012; Mell et al., 2014). Experimental work indicates that the distribution of eDNA length acquired is skewed towards short fragments, with a third of transformation events less than 1kb, a median around 2-6kb and range extending up to ∼50kb (Croucher et al., 2012). This appears to be an evolved state in *Streptococcus pneumoniae*, as the dedicated system cleaves eDNA into smaller fragments before recombination takes place (Claverys et al., 2009). Some studies have reported a larger median and range for transfer sizes (Hiller et al., 2019). Given that loci are around 1kb, with short intergenic regions, this represents the potential for several genes to be transferred in a single LGT event (Mira et al., 2001; Moran, 2002). There are several potential reasons for focus on small genomic pieces in LGT recombination. Cleavage of eDNA into smaller sequences increases the likelihood of homologous recombination, while the acquisition of long sequences can be associated with loss of genetic information (Croucher et al., 2012) and can potentially disrupt regulatory and physiological networks (Baltrus, 2013). It has also been suggested that the small size of recombination fragments is a mechanism for preventing the spread of mobile genetic elements (Croucher et al., 2016). On the other hand, gram-negative bacteria do not cleave eDNA on import, but their ability to acquire eDNA sequences >50kb is limited by physical constraints (Mell and Redfield, 2014). The high variability of LGT size suggest that there is flexibility and the potential for evolutionary change. But there is no evidence that larger genome size is accompanied by a higher recombination length. To our knowledge bacteria do not load large pieces of chromosome via LGT (i.e. >10% of a genome), although in principle it should be possible for them to do so.

As for purely asexual populations (Haigh, 1978), the strength of selection plays a critical role in determining the rate of mutation accumulation, with regions of the genome under strong selection accumulating mutations at a low rate (Fig. 5-6). The ratchet effect is mainly observed in the accessory genome, with mutations accumulating preferentially in loci under weak selection (Fig. 5-6). Our model predicts that genome size expansion can occur in populations under strong purifying selection (e.g. due to a larger effective population size). Strong selection decreases the rate of genetic information loss, allowing the acquisition of new genetic content without an attendant increase in mutation fixation. This prediction is in agreement with the positive correlation observed between genome size and dN/dS in bacteria (Novitchkov et al., 2008; Bobay and Ochman, 2018). However, organisms under similar selective pressures often display a broad range of genome sizes (Novitchkov et al., 2008), indicating that other factors, including mutation rate and LGT, have a strong impact on prokaryotic genome size. Under high mutation rate and weak selective pressure, genome size expansion is disfavored.

Eukaryotes, including simple unicellular organisms, typically possess much larger genomes than prokaryotes, as noted earlier (Koonin, 2009; Elliot and Gregory, 2015). Eukaryotic genome size expansion was favored by the acquisition of an endosymbiont, which evolved into the mitochondrion. This released bioenergetic constraints on cell size and allowed the evolution of genetic and morphological complexity (Lane, 2014; Lane, 2020). The endosymbionts underwent gene loss, a frequently observed process in extant endosymbiotic relationships (López-Madrigal and Rosario, 2017) and transferred multiple genes to the host, enriching the host’s genome size with genes of proto-mitochondrial origin (Timmis et al., 2004; Martin et al., 2015). The acquisition of endosymbiotic DNA is also thought to have allowed the spread of mobile genetic elements in the host cell’s genome, contributing to the increase in genome size and likely increasing the mutation rate (Timmis et al., 2004; Martin and Koonin, 2006; Rogozin et al., 2012). The loss of energetic constraints on genome size probably also facilitated gene and even whole genome duplications, leading to several thousand new gene families in LECA (Koonin, 2004), as well as lower selective pressure for gene loss after LGT (Szollosi, Derenyi and Vellai, 2006).

Such a large genome brought the first eukaryotes under the threat of mutational accumulation, creating the need for stronger purifying selection in order to keep the expanded genetic content free from mutations. Our results offer a possible explanation of why this process drove the transition from LGT to meiotic recombination at the origin of sex. In small prokaryotic genomes, LGT provides sufficient benefits to maintain genome integrity, without incurring the multiple costs associated with sexual reproduction. But LGT fails to prevent the accumulation of deleterious mutations in larger genomes, promoting the loss of genetic information and therefore constraining genome size. Our model shows that genome size expansion is only possible through a proportional increase in recombination length. We considered a recombination length *L* = 0.2*g*, which is equivalent to 500 genes for a species with genome size of 2,500 genes – two orders of magnitude above the average estimated eDNA length in extant bacteria (Croucher et al., 2012). Recombination events of this magnitude are unknown among prokaryotes, possibly because of physical constraints on eDNA acquisition. Limiting factors likely include the restricted length of eDNA, uptake kinetics and the absence of an alignment mechanism for large eDNA strands (Thomas and Nielsen, 2005; Baltrus, 2013; Croucher et al., 2016).

The requirement for a longer recombination length *L* cannot be achieved by LGT, which must therefore have failed to maintain a mutation-free genome, generating a strong selective pressure towards the evolution of a new mechanism of inheritance with the loss of energetic constraints on genome size. However, this magnitude of *L* is easily achievable via meiotic sex. The transition from LGT to meiotic sex involves the evolution of cell fusion, the transition from circular to linear chromosomes, whole-chromosome alignment and homologous recombination (Lane, 2011; Goodenough and Heitman, 2014). We have not explicitly modelled the details of this process or considered the order in which these factors arose. Nonetheless, our results show that eukaryotes had to increase the magnitude of recombination length beyond the limits of LGT in order to permit the expansion in genetic complexity without the attendant increase in mutational burden. Eukaryotes had to abandon LGT in order to increase recombination length and maintain a large genome. Sex was forced upon us.

## CONCLUSION

The benefits of LGT in maintaining genome integrity decline rapidly with genome size, making large genomes vulnerable to the accumulation of mutations. This effect constrains genome size in prokaryotes, and becomes even more severe with small population sizes and high mutation rates. These constraints can be partially overcome by increases in LGT rate and recombination length (Fig 3-4). But only recombination across the whole genome can wholly overcome these constraints. With the massive genome expansion at the origin of eukaryotes, the evolution of meiosis allowed homologous recombination across the whole genome, and not only across a limited region spanning little more than a few loci, as in LGT. The endosymbiosis that gave rise to the first eukaryotes led to the frequent transfer of genes from the endosymbiont to the host, resulting in a large expansion in genome size, likely coupled to high mutation rates. Our model shows that these conditions wrought the failure of LGT in preventing Muller’s ratchet. The resulting selective pressure promoted the evolution of sexual cell fusion and meiosis, maximizing recombination length and protecting eukaryotic genomes from excessive mutational burden. LGT in prokaryotes gave way to meiotic sex in eukaryotes because only sex can sustain the expansion in genome size that underpins all eukaryotic complexity.

---

**Table 1.** Parameters and variables.

|  |  |
| --- | --- |
| $N$ | *population size* |
| $\mu$ | *mutation rate per bp per generation* |
| $g$ | *genome size (number of loci)* |
| $U$ | *genome-wide mutation rate* |
| $s$ | *strength of selection against deleterious mutations* |
| $\lambda$ | *LGT rate* |
| $L$ | *eDNA length (number of loci)* |

## REFERENCES

Ambur, O. H., Engelstädter, J., Johnsen, P. J., Miller, E., and Rozen, D. E. 2016 Steady at the wheel: conservative sex and the benefits of bacterial horizontal gene transfer. Philos. Trans. R Soc. Lond. B Biol. Sci. 371:20150528.

Baltrus, D. A. 2013 Exploring the costs of horizontal gene transfer. Trends Ecol. Evol. 28: 489–495.

Barton, N. H., and Otto S. P. 2005 Evolution of recombination due to random drift. Genetics 169: 2353–2370.

Bell, G. 1982 The masterpiece of nature: the evolution and genetics of sexuality. Berkeley, CA: University of California Press.

Bernstein, H., and Bernstein, C. 2013 Evolutionary origin and adaptive function of meiosis. Meiosis, 1.

Bobay, L., and Ochman, H. 2018 Factors driving effective population size and pan-genome evolution in bacteria. BMC Evolutionary Biology 18: 153.

Carr, M., Leadbeater, B. S., and Baldauf, S. L. 2010 Conserved meiotic genes point to sex in the choanoflagellates. J. Eukaryot. Microbiol. 57: 56–62.

Claverys, J. P., Martin, B., and Polard, P. 2009 The genetic transformation machinery: composition, localization, and mechanism. FEMS Microbiol. Rev. 33: 643–656.

Croucher, N. J., Mostowy, R., Wymant, C., Turner, P., Bentley, S. D., and Fraser, C. 2016 Horizontal DNA transfer mechanisms of bacteria as weapons of intragenomic conflict. PLoS Biol. 14: e1002394.

Croucher, N. J., Harris, S. R., Barquist, L., Parkhill, J., and Bentley, S. D. 2012 A high-resolution view of genome-wide pneumococcal transformation. PLoS Pathog. 8: e1002745.

Elliot, T. A., and Gregory, T. R. 2015 What’s in a genome? The C-value enigma and the evolution of eukaryotic genome content. Phil. Trans. R. Soc. B 370: 20140331.

Felsenstein, J., and Yokoyama, S. 1976 The evolutionary advantage of recombination. II. Individual selection for recombination. Genetics 83: 845–859

Fuchsman, C. A., Collins, R. E., Rocap G., and Brazelton W. J. 2017 Effect of the environment on horizontal gene transfer between bacteria and Archaea. PeerJ 5: e3865.

Gandon, S., and Otto, S. P. 2007 The evolution of sex and recombination in response to abiotic or coevolutionary fluctuations in epistasis. Genetics 175: 1835–1853.

Goodenough, U., and Heitman, J. 2014 Origins of eukaryotic sexual reproduction. Cold Spring Harbor Perspect. Biol. 6, a016154.

Gordo, I., and Charlesworth, B. 2000 The degeneration of asexual haploid populations and the speed of Muller’s ratchet. Genetics 154: 1379–1387.

Hamilton, W. D. 1980 Sex vs. non-sex vs. parasite. Oikos 35: 282–290.

Haigh, J. 1978 The accumulation of deleterious genes in a population—Muller’s ratchet. Theor. Pop. Biol. 14: 251–267.

Hao, W., and Golding, G. B. 2006 The fate of laterally transferred genes: life in the fast lane to adaptation or death. Genome Res. 16: 636–643.

Hiller, N. L., Ahmed, A., Powell, E., Martin, D. P., Eutsey R., et al. 2010 Generation of genic diversity among *Streptococcus pneumoniae* strains via horizontal gene transfer during a chronic polyclonal pediatric infection. PLoS Pathog. 6: e1001108.

Hofstatter, P. G., Brown, M. W., and Lahr, D. J. 2018 Comparative genomics supports sex and meiosis in diverse Amoebozoa. Genome Biol. Evol. 10: 3118–3128.

Jain, R., Riviera, M. C., Moore, J. E., and Lake, J. A. 2003 Horizontal gene transfer accelerates genome innovation and evolution. Mol. Biol. Evol. 20: 1598–1602.

Johnston, C., Martin, B., Fichant, G., Polard, P., and Claverys, J-P. 2014 Bacterial transformation: distribution, shared mechanisms and divergent control. Nat. Rev. Microbiol. 12: 181–96.

Jokela, J., Dybdahl, M. F., and Lively, C. M. 2009 The maintenance of sex, clonal dynamics, and host-parasite coevolution in a mixed population of sexual and asexual snails. Am. Nat. 174: S43–S53.

Koonin, E. V. 2009 Evolution of genome architecture. Int. J. Biochem. Cell Biol., 41: 298–306.

Koonin, E. V., Fedorova N., D., Jackson J. D., Jacobs A., R., Krylov D., M., et al. 2004 A comprehensive evolutionary classification of proteins encoded in complete eukaryote genomes. Genome Biol. 5: R7.

Lahr, D. J., Parfrey, L. W., Mitchell, E. A., Katz, L. A., and Lara, E. 2011 The chastity of amoebae: re-evaluating evidence for sex in amoeboid organisms. Proc. R. Soc. B Biol. Sci. 278: 2081–2090.

Lane N. 2020 How energy flow shapes cell evolution. Curr. Biol. (in press).

Lane, N. 2014 Bioenergetic constraints on the evolution of complex life. Cold Spring Harbor Perspect. Biol. 6: a015982.

Lane, N. 2011 Energetics and genetics across the prokaryote-eukaryote divide. Biol. Direct 6: 35.

Lane, N., and Martin, W. 2010 The energetics of genome complexity. Nature 467: 929–934.

Lapierre, P., and Gogarten, J. P. 2009 Estimating the size of the bacterial pan-genome. Trends Genet. 25: 107–110.

Lenormand, T., and Otto S. P. 2000 The evolution of recombination in a heterogeneous environment. Genetics 156: 423–438.

Levin, B.R., and Cornejo, O. E. 2009 The population and evolutionary dynamics of homologous gene recombination in bacterial populations. PLoS Genet. 5: e1000601.

Lin, Z., Kong, H., Nei, M., and Ma, H. 2006 Origins and evolution of the *RecA/RAD51* gene family: Evidence for ancient gene duplication and endosymbiotic gene transfer. Proc. Nat. Aca. Sci. USA 103: 10328–10333.

López-Madrigal S., and Rosario G. 2017 Et tu, Brute? Not even intracellular mutualistic symbionts escape horizontal gene transfer. Genes 8: 247.

Malik, S. B., Pightling, A. W., Stefaniak, L. M., Schurko, A. M., and Logsdon Jr, J. M. 2008 An expanded inventory of conserved meiotic genes provides evidence for sex in Trichomonas vaginalis. PloS one 3: 8.

Marri, P. R., Hao., W., and Golding, G. B. 2006 Gene gain and gene loss in *Streptococcus*: is it driven by habitat? Mol. Biol. Evol. 23: 2379–2391.

Marri, P. R., Hao., W., and Golding, G. B. 2007 The role of laterally transferred genes in adaptive evolution BMC Evol. Biol. 7: S8.

Martin, W., and Koonin, E. V. 2006 Introns and the origin of nucleus-cytosol compartmentalization. Nature 440: 41–45.

Martin, W., Garg, S., and Zimorski, V. 2015 Endosymbiotic theories for eukaryote origin. Philos. Trans. R. Soc. Lond. B. Biol. Sci. 370: 20140330.

Mell., J. C., and Redfield, R. 2014 Natural competence and the evolution of DNA uptake specificity. J Bacteriol. 196: 1471–1483.

Mell, J. C., Lee, J. Y., Firme, M., Sinha, S., and Redfield, R. J. 2014 Extensive cotransformation of natural variation into chromosomes of naturally competent *Haemophilus influenzae*. G3 4: 717–731.

Mirzaghaderi, G., and Horandl, E. 2016 The evolution of meiotic sex and its alternatives. Proc. R. Soc. B 283: 20161221.

Mira, A., Ochman, H., and Moran, N. A. 2001 Deletional bias and the evolution of bacterial genomes. Trends Genet. 17: 589–596.

Moran, N. A. 2002 Microbial minimalism: genome reduction in bacterial pathogens. Cell 108: 583–586.

Muller, H. J. 1964 The relation of recombination to mutational advance. Mutat. Res. Mol. Mech. Mutagen. 1: 2–9.

Müller, M., Mentel, M., van Hellemond, J. J., Henze, K., Woehle, C., et al. 2012 Biochemistry and evolution of anaerobic energy metabolism in eukaryotes. Microbiol. Mol. Biol. Rev. 76: 444–495.

Novichkov, P. S., Wolf, Y. I., Dubchack, I., and Koonin, E. V. 2009 Trends in prokaryotic evolution revealed by comparison of closely related bacterial and archaeal genomes. Journal of Bacteriology 191: 65–73.

Nowell, R. W., Green, S., and Sharp, P. M. 2014 The extent of genome flux and its role in the differentiation of bacterial lineages. Genome Biol. Evol. 6: 1514–1529.

Ochman, H., Lawrence, J. G., and Groisman, E. A. 2000 Lateral gene transfer and the nature of bacterial innovation. Nature 405: 299–304.

Otto, S. P., and Lenormand, T. 2002 Resolving the paradox of sex and recombination. Nat. Rev. Genet. 3: 252–261.

Otto, S. P. 2009 The evolutionary enigma of sex. Am. Nat 174: S1–S14.

Pylkov, K. V., Zhivotovsky, L. A., and Feldman, M. W. 1998 Migration versus mutation in the evolution of recombination under multi-locus selection. Genet. Res. 71: 247–256.

Ramesh, M. A., Malik, S. B., and Logsdon Jr, J. M. 2005 A phylogenomic inventory of meiotic genes: evidence for sex in Giardia and an early eukaryotic origin of meiosis. Curr. Biol. 15: 185–191.

Redfield, R. J. 1988 Evolution of bacterial transformation: is sex with dead cells ever better than no sex at all? Genetics 119: 213–221.

Redfield, R. J., Schrag, M. R., and Dean, A. M. 1997 The evolution of bacterial transformation: sex with poor relations. Genetics, 146: 27–38.

Rogozin, I. B., Carmel, L., Csuros, M., and Koonin, E. V. 2012 Origin and evolution of spliceosomal introns. Biol. Direct 7:11.

Seitz, E. M., Brockman, J. P., Sandler, S. J., Clark, A. J., and Kowalczykowski, S. C. 1998 RadA protein is an archaeal RecA protein homolog that catalyzes DNA strand exchange. Genes Dev. 12: 1248–1253.

Sela, I., Wolf, Y. I., and Koonin, E. V. 2016 Theory of prokaryotic genome evolution. Proc. Natl. Acad. Sci. USA 113: 11399–11407.

Schurko, A. M., Logsdon, J. M. 2008 Using a meiosis detection toolkit to investigate ancient asexual ‘scandals’ and the evolution of sex. BioEssays 30: 579–589.

Speijer, D., Lukes, J., Elias, M. 2015 Sex is a ubiquitous, ancient, and inherent attribute of eukaryotic life. Proc. Natl. Acad. Sci. USA 112: 8827–8834.

Szollosi, G. J., Derenyi, I., Vellai, T. 2006 The maintenance of sex in bacteria is ensured by its potential to reload genes. Genetics 174: 2173–2180

Takeuchi, N., Kaneko, K., and Koonin, E. V. 2014 Horizontal gene transfer can rescue prokaryotes from Muller’s ratchet: benefit of DNA from dead cells and population subdivision. G3 4: 325–339.

Thomas, C. M., and Nielsen, K. M. 2005 Mechanisms of, and barriers to, horizontal gene transfer between bacteria. Nature Rev. Microbiol. 3: 711–21.

Timmis, J. N., Ayliffe, M. A., Huang, C. Y., and Martin, W. 2004 Endosymbiotic gene transfer: organelle genomes forge eukaryotic chromosomes. Nature Rev. Genet. 5: 123–135.

Vos, M. 2009 Why do bacteria engage in homologous recombination? Trends Microbiol. 17: 226–232.

Vos, M., Hesselman, M. C., te Beek, T. A., van Passel, M. W., and Eyre-Walker, A. 2015 Rates of lateral gene transfer in prokaryotes: high but why? Trends Microbiol. 23: 598–605.

Williams, T. A., Foster, P. G., Cox, C. J., and Embley, T. M. 2013 An archaeal origin of eukaryotes supports only two primary domains of life. Nature 504: 231–236.

Wylie, C. S., Trout, A. D., Kessler, D. A., and Levine, H. 2010 Optimal strategy for competence differentiation in bacteria. PLoS Genet. 6: e1001108.

Zaremba-Niedzwiedzka, K., Caceres, E. F., Saw, J. H., Bäckström, D., Juzokaite, L., et al. 2017 Asgard archaea illuminate the origin of eukaryotic cellular complexity. Nature 541: 353–358.

